# Automated single-cell omics end-to-end framework with data-driven batch inference

**DOI:** 10.1101/2023.11.01.564815

**Authors:** Yuan Wang, William Thistlethwaite, Alicja Tadych, Frederique Ruf-Zamojski, Daniel J Bernard, Antonio Cappuccio, Elena Zaslavsky, Xi Chen, Stuart C. Sealfon, Olga G. Troyanskaya

## Abstract

To facilitate single-cell multi-omics analysis and improve reproducibility, we present SPEEDI (Single-cell Pipeline for End to End Data Integration), a fully automated end-to-end framework for batch inference, data integration, and cell type labeling. SPEEDI introduces data-driven batch inference and transforms the often heterogeneous data matrices obtained from different samples into a uniformly annotated and integrated dataset. Without requiring user input, it automatically selects parameters and executes pre-processing, sample integration, and cell type mapping. It can also perform downstream analyses of differential signals between treatment conditions and gene functional modules. SPEEDI’s data-driven batch inference method works with widely used integration and cell-typing tools. By developing data-driven batch inference, providing full end-to-end automation, and eliminating parameter selection, SPEEDI improves reproducibility and lowers the barrier to obtaining biological insight from these valuable single-cell datasets. The SPEEDI interactive web application can be accessed at https://speedi.princeton.edu/.

## Introduction

Single-cell sequencing technologies provide access to the molecular environment of individual cells in different tissues, species, and conditions. Recent advances in multi-modal assays have expanded the ability to study the behavior of complex cellular systems at single-cell resolution. Single-cell methods can provide insight not achievable with traditional bulk assay technology. While interest in single-cell technologies by clinical and translational researchers has been rapidly growing, analysis of these datasets is not routine and requires a high level of bioinformatics expertise for proper interpretation.

Recovering a robust single-cell landscape and identifying differential patterns between cell types or conditions entails interrogating datasets from multiple samples. These analyses currently require the use of multiple software tools as well as Python or R scripting. The need for familiarity with rapidly evolving single-cell analysis methods creates a bottleneck for many biology and biomedical research groups in obtaining robust interpretations of publicly accessible or laboratory generated single-cell datasets. Importantly, the manual parameter optimization needed for the use of current tools may lead to different conclusions from the same data.

A key methodological challenge is that variations in the time of sample processing, sequencing depth, or unknown technical factors result in batch effects that compromise integration, analysis, and interpretation of single-cell studies. Notably, exploration of single-cell data (for example on UMAP plots or PCA plots) in studies of multiple samples often shows evidence of sample groups that do not represent either biological factors (e.g. a contrast being studied) or any recorded experimental batch factors. While existing single-cell integration methods and benchmarking studies all assume that batch information is provided, accurate, and complete,^1,2,3,4,5,6^ identification of batches can be a particularly difficult problem as the origin of many batch effects is unknown. An additional issue is that experimentally-recorded batch factors may not in fact cause any sample grouping--i.e. they do not actually cause batch effects. Including batch labels that do not have significant batch effects can severely reduce the statistical power of the analysis and obscure biological differences between the experimental contrasts under study (Figure 1A).

**Figure 1.**
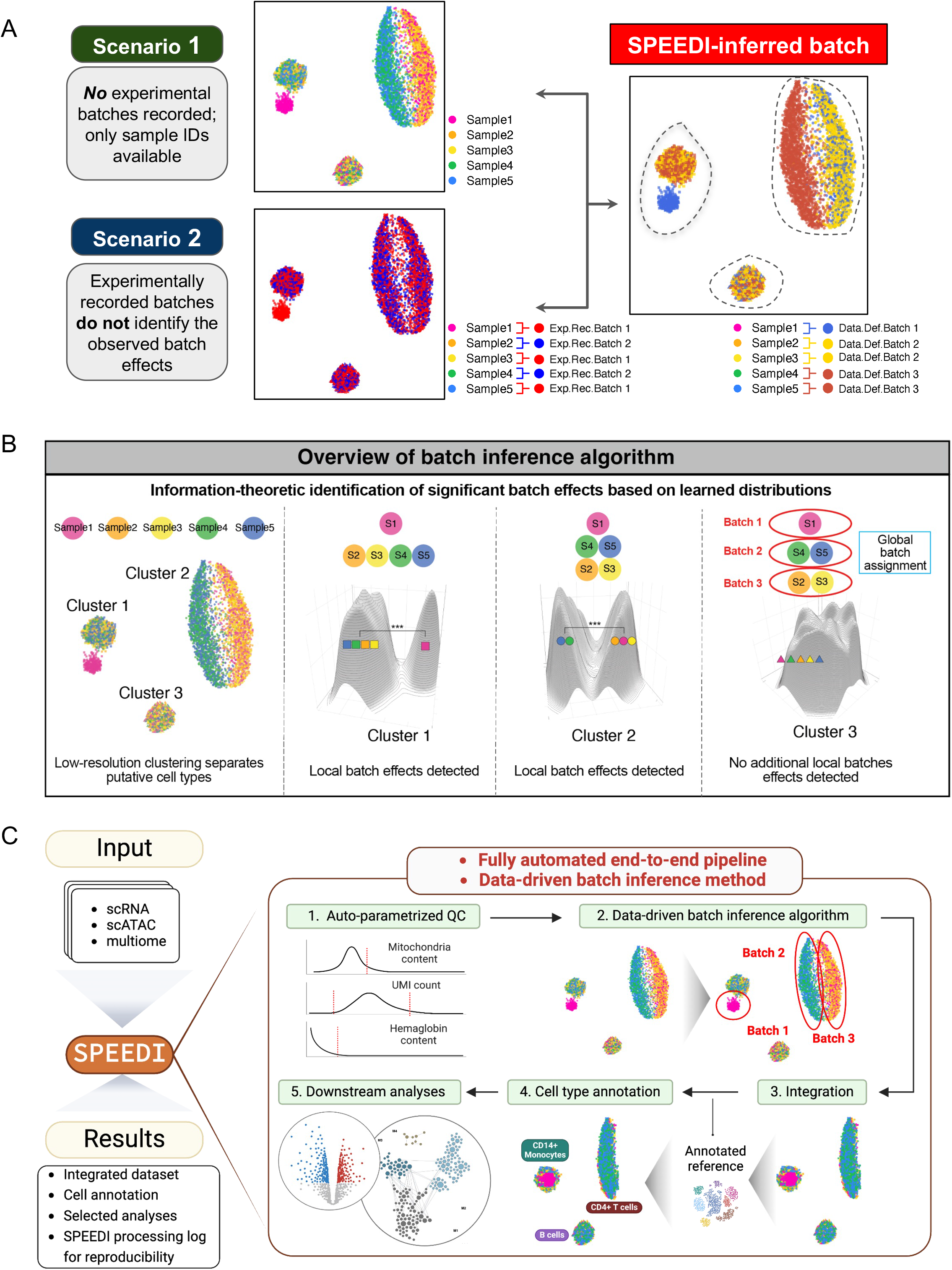
Schematic illustrations of potential batch reporting scenarios and SPEEDI’s solution (see also Table S1). (**A**) Different batch effect scenarios. Single-cell data commonly shows non-biological batch effects where the data from groups of samples shows distinct patterns. These batch effects may result from known experimentally recorded factors (which we refer to as experimentally-recorded batches). However, the data often shows non-biological batch effects that are not annotated to any known experimental factor. Whatever the cause of batch effects, they need identification and correction to improve the rigor of detection of true biological effects. The limitations of experimentally-recorded batch identification are illustrated schematically by showing a single-cell dataset of individual samples from 5 subjects comprising three cell types in which batch effects in the data are observed. (Data coming from the same cell type are enclosed within the dashed outlines in the SPEEDI inferred batch panel.) The observed batch effects are not adequately labeled in scenarios in which no batch information is provided (scenario 1) or the experimentally recorded batches do not correctly identify any or all the batch effects present (scenario 2). A data-driven method to provide accurate batch identification, whether due to experimentally-recorded factors or due to unknown factors, is needed. (**B**) SPEEDI uses an information-theoretic approach to quantify sample distributions and identify local and global batch effects based on learned distributions. The algorithm starts with a low-resolution clustering that separates putative cell types. It then iterates through the clusters to determine if a group of samples is significantly different from the rest of the samples and assigns batch labels locally. In cases where a subset of samples within the significant batch has already been assigned a batch label in the previous iteration, the framework further divides the significant batch by giving the previously unassigned samples a new batch label. This process is repeated until all local batches are stable, which constitutes the final global batch assignment. (**C**) SPEEDI provides a one-step, fully automated multiple sample single-cell analysis pipeline that does not require any parameter selection by the user and includes a data-driven batch inference method to improve the quality of integration. The framework takes the CellRanger output and implements a workflow comprising quality control with automated parameter selection (Step 1), application of the data-driven batch inference method (Step 2), data integration (Step 3), cell type annotation (Step 4), and optional downstream differential and pathway analyses (Step 5). SPEEDI returns an annotated, integrated data matrix for single-cell RNA-seq and/or for single-cell ATAC-seq, as well as selected analyses.

To clarify these issues, we define two different types of non-exclusive “batches”: 1. Experimentally-recorded batch: groups of samples sharing a common experimental factor (such as library processing groups). Experimentally-recorded batches may or may not be associated with actual differences seen between the data obtained from samples in these groups. This is the current use of the term “batch.” 2. Data-defined batch: Grouping of data from sample subsets that do not represent biological factors (e.g. a contrast being studied). Data-defined batches may correspond to some experimentally recorded technical factors, but often this is not the case. Whether the source of data-defined batch effects is unknown or can be attributed to known experimental factors, these effects need to be corrected (and experimental factors that do not in fact cause batch effects need to be ignored) for rigorous interpretation of the data. Therefore, a method that identifies batches purely in a data-driven manner is needed to identify and correct batch effects whatever their origin.

We address these barriers to analysis of single-cell datasets with the SPEEDI (Single-cell Pipeline for End-to-End Data Integration) framework. SPEEDI introduces the first automated data-driven batch inference method, overcoming the problem of unknown or under-specified batch effects (**Figure 1B**). SPEEDI is a fully automated end-to-end QC, data-driven batch identification, data integration, and cell-type labeling pipeline that does not require any manual parameter selection or pipeline assembly (**Figure 1C, Table S1**). SPEEDI currently works with scRNA, scATAC, or sc multiome datasets and matches or exceeds benchmarking against existing manual, piecemeal methods. Unlike these other methods that also require a high level of bioinformatics expertise, SPEEDI is available both in a web interface and an R package, making it accessible to a wide variety of researchers. SPEEDI’s full automation improves reproducibility and robustness of analyses, and democratizes the interpretation of single-cell omics datasets.

## Results

### SPEEDI framework

Accounting for batch effects of unknown origin in single-cell data is critical in correcting for non-biological artifacts that compromise data analysis and interpretation. To address this problem, we developed a data-driven batch identification method based on information-theoretic principles (**Figure 1B**; see **STAR Methods**).

This novel data-driven batch identification method, combined with automated parameter selection, enabled us to build SPEEDI, a comprehensive, automated end-to-end single-cell data analysis workflow (**Figure 1C**). SPEEDI automates quality control and low-quality cell filtering, batch label inference, multi-sample integration, cell type annotation, and a variety of user-selected downstream analyses, while seamlessly integrating the functionality of existing single-cell analysis packages, such as Seurat and ArchR. Low-quality cell filtering is automated through distribution-based thresholding applied to multiple quality control criteria relevant for the specific data type (see **STAR Methods**).

In the default general user implementation, following cell type labeling, SPEEDI incorporates public methods to integrate samples and annotate cell types.^2,4^ SPEEDI further refines cell type annotation by introducing a majority-based voting algorithm (see **STAR Methods**). Users can either choose among the annotation references provided with SPEEDI or can use their own reference. For advanced users, SPEEDI can use any parametric integration method that accepts an input list of datasets and batch labels, such as Scanorama or Harmony,^3,4^ and accommodates other single-cell annotation tools, such as SingleR.^7^

### Validation of data-driven sample batch inference method and SPEEDI framework

We evaluated the SPEEDI framework using four different real-world case studies that included different data types. We first quantified the performance of the batch inference method applied to a public human lung scRNA-seq dataset using the study authors’ curated and published cell type annotation as the reference. Second, we computationally inferred batches from a human PBMC scRNA-seq dataset and compared the cell type annotations to those obtained from using sample ID information. Third, we showed that SPEEDI is robust across different data modalities by analyzing a mouse same-cell multiome dataset. Finally, we demonstrate the utility of the downstream analysis tools in SPEEDI by evaluating human PBMC scRNA-seq data from a longitudinal SARS-CoV-2 study.

### Benchmarking the SPEEDI batch inference method

The human healthy lung single-cell expression data, originally collected by Braga et al.,^8^ includes three diverse datasets across different sequencing technologies. Two of the datasets were generated by 10X Chromium and the other one was generated by Drop-seq. We used the cell type annotation of the three datasets comprising 17 cell types and 16 samples in total reported by Luecken et al. as an expert analysis standard for benchmarking.^5^

The heterogeneity of samples resulted in the presence of strong batch effects in the data that made integration particularly challenging. Other than intrinsic inter-individual variations, the different sequencing protocols and laboratories generating the data resulted in highly variable cell type compositions between samples (**Table S2**). We compared the results obtained using the following different batch labels: i. batch label = each sample ID ii. batch label = each dataset ID (i.e. data on sets of samples that are obtained from different experiments that are processed separately) iii. batch label = SPEEDI data-defined batches. We compared the results obtained with each batch labeling approach using four different integration methods: Seurat CCA, Seurat RPCA, Harmony, and Scanorama. We then applied four different scoring metrics that assess preservation of cell type information and removal of dataset-specific variations to provide a comprehensive comparison of the batch labeling methods (**Figure 2**; see **STAR Methods**).^5^ SPEEDI batch labeling led to a significant performance improvement in nearly all comparisons with the two Seurat and Harmony integrations. The improvement of SPEEDI using Scanorama, which provides lower quality results for most metrics, was more modest. Overall, the benchmarking study indicates that the SPEEDI batch inference algorithm significantly boosts integration performance.

**Figure 2.**
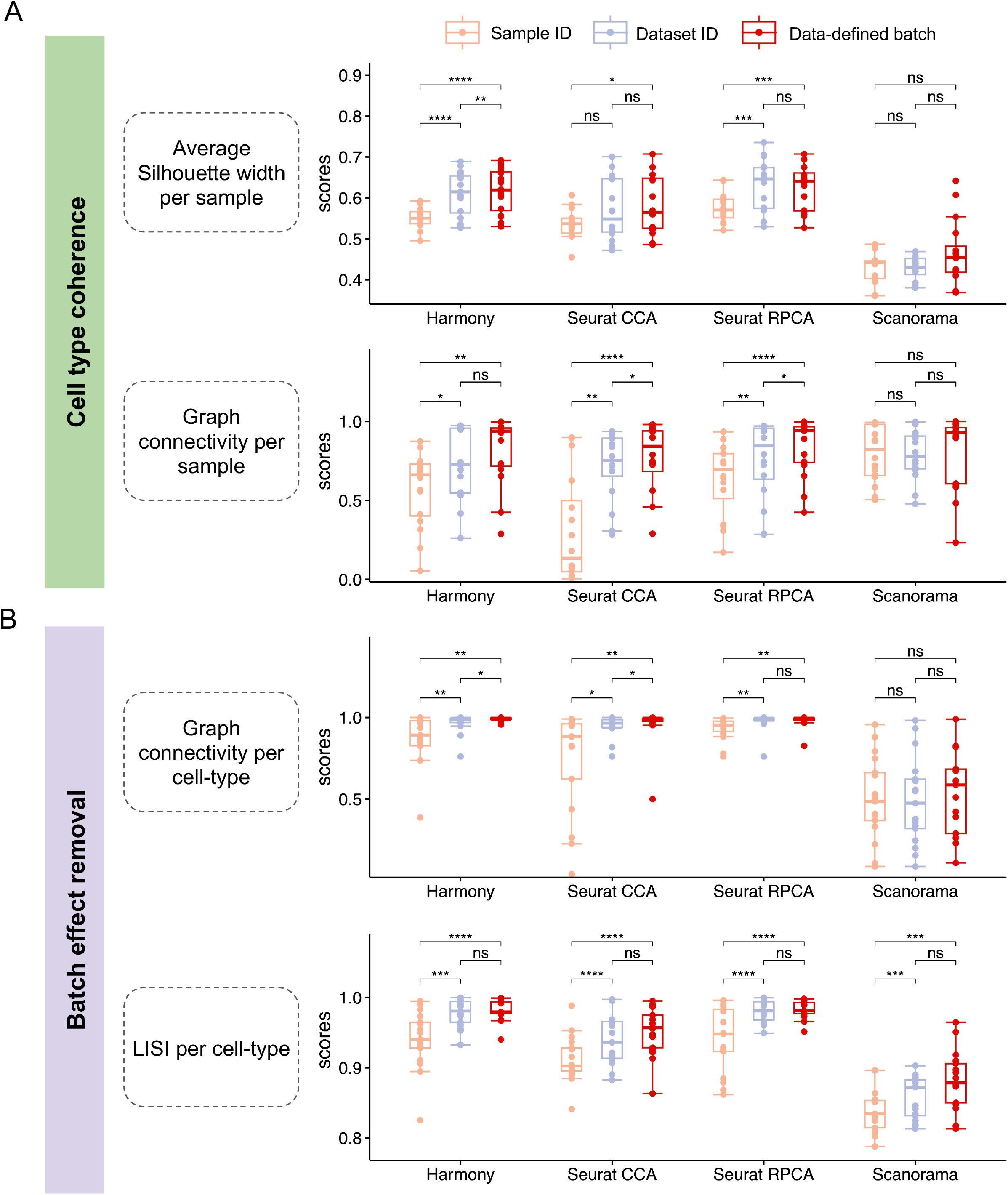
Benchmarking batch labeling strategies (see also Table S2). In a public atlas-level human lung scRNA-seq dataset comprising data from three different datasets, batches were labeled using either the sample ID, the dataset ID, or the SPEEDI data-driven method. The three types of batch labels were used as input for four different batch correction and data integration packages (Harmony, Seurat CCA, Seurat RPCA, Scanorama). The results of the integration with different batch labeling approaches were compared using metrics for cell type coherence and for batch removal. (**A**) Evaluation of SPEEDI batch inference in preserving cell type coherence among distinct cell types after integration. Each dot represents a sample score for a total of 16 samples. Each score quantifies the effectiveness in preserving the integrity of different cell types within that sample. A higher score indicates that the cells of a particular cell type are more distinctly separated from cells of other types after integration. Pairwise nonparametric Wilcoxon rank sum tests were performed, and p-values were Bonferroni-adjusted. (**B**) Evaluation of SPEEDI batch inference in eliminating batch variants among samples after integration. East dot represents a cell type for a total of 17 types. Each score represents how effectively batches are mitigated within the associated cell type. A higher score indicates that cells of the same type, regardless of their sample origin, are better inter-mixed after integration. Pairwise nonparametric Wilcoxon rank sum tests were performed, and p-values were Bonferroni-adjusted.

### Application of SPEEDI to heterogeneous single-cell RNA-seq data

To further evaluate the reliability and robustness of the SPEEDI batch inference method and overall framework, we applied SPEEDI to a 20 subject scRNA-seq dataset generated in our laboratory for a study of the responses in peripheral blood mononuclear cells (PBMC) to *S. aureus* infection.^9^ Before batch correction, a number of UMAP clusters were distinguished by sample, a pattern consistent with the presence of severe batch effects (see **Figure 3A** for T/NK lymphocyte subpopulations and **Figure S1** for all PBMC cell types). SPEEDI data-inferred batch labels led to much better batch correction compared with batch labels obtained from sample IDs (**Figure 3B** and **Figure S2**). For example, SPEEDI automated batch labeling corrected the fragmentation of CD14 monocytes and Treg cells into different clusters as seen when using sample ID batch correction (**Figure S3**). Furthermore, when using sample ID labels for batch correction, CD8+ memory cells were more broadly distributed, intermingled with other T cells, and showed less overlap with CD8A versus using data-defined batch labels for batch correction (Figure S4). Expression plots for canonical marker genes (NKG7 for NK cells, SELL for CD4 naive cells, TNFRSF4 for CD4 memory cells, and FOXP3 for Treg cells) confirmed SPEEDI inferred cell types (Figure S5). The SPEEDI method identified a complex pattern of 12 data-defined batches that varied in size from 1 to 5 samples. (**Figure 3C**). Application of the four different batch correction quality scoring metrics used in the lung dataset above showed that using SPEEDI batch identification performed better than using sample IDs for three of the metrics and comparably for the remaining metric (**Figure 3D**). These results demonstrate that, in comparison with sample ID labels, using SPEEDI data-defined batch labels improves the accuracy of dataset integration and cell type annotation.

**Figure 3.**
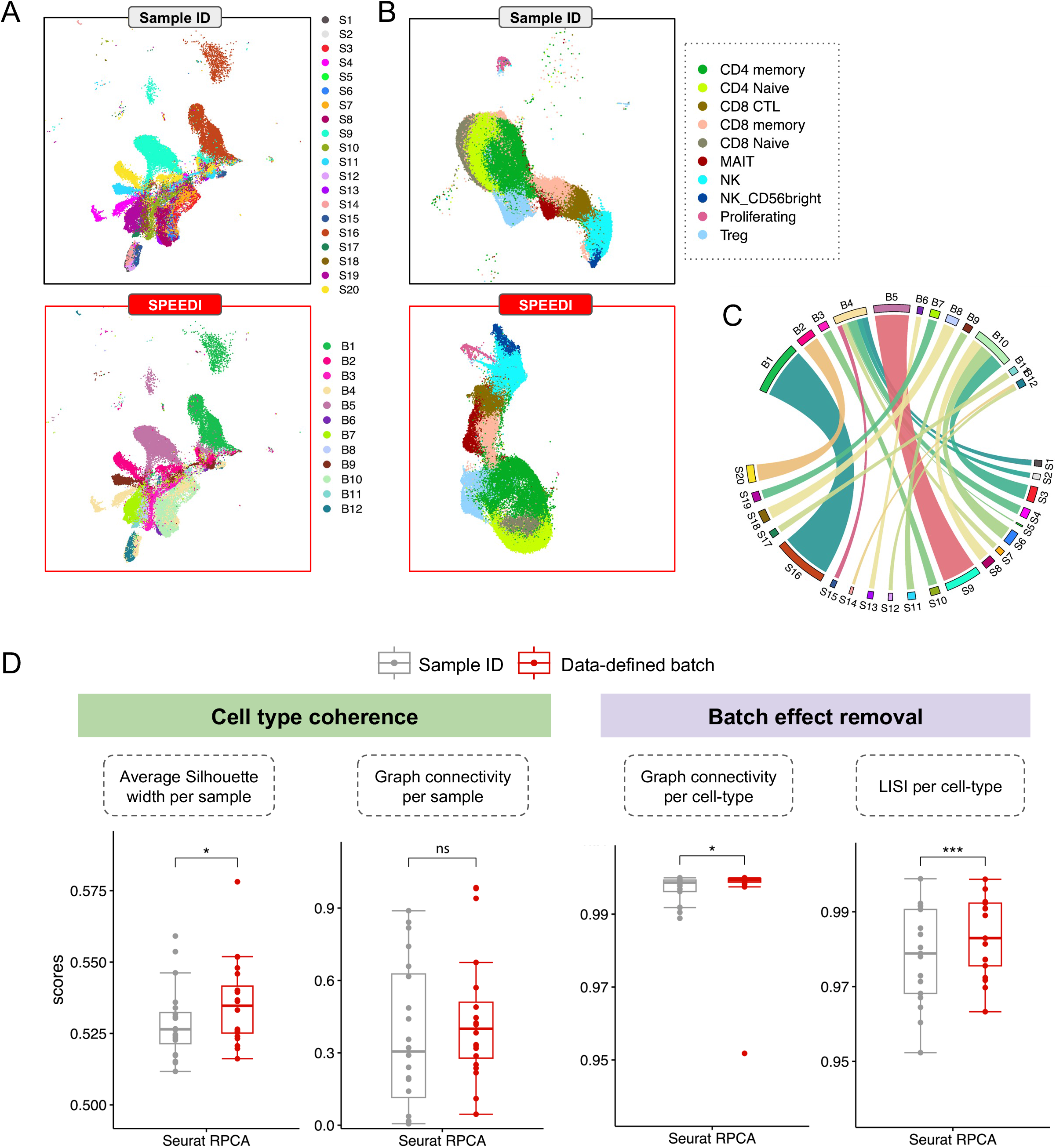
SPEEDI facilitates integrative cell type identification from scRNA-seq data. (See also Figures S1-5) (**A**) We applied SPEEDI to a human peripheral blood mononuclear cell (PBMC) scRNA-seq study with 20 subjects. Batches were identified using either the automated data-driven batch inference method implemented in SPEEDI (n = 12) or using sample IDs (n = 20). The UMAPs show the T/NK cell population before integration (see Figure S1 for UMAPs with all major PBMC cell types). (**B**) The UMAPs of the T/NK cell population after integration with both batch labeling strategies are shown (see Figure S2 for UMAPs with all major PBMC cell types). (**C**) Correspondence between the data-defined batch labels and sample IDs. (**D**) Score for cell type coherence. For cell type coherence measures, each dot represents a sample score, and each score quantifies the effectiveness in preserving the integrity of different cell types within that sample. Pairwise nonparametric Wilcoxon rank sum tests were performed. Scores for batch removal. For batch effect removal metrics, each dot represents a cell type score, and each score represents how effectively batches are mitigated within the associated cell type. Pairwise nonparametric Wilcoxon rank sum tests were performed. *p<0.05, *** p<0.001, n.s. Not-significant (p>0.05). Bonferroni corrected t-test.

### Application of SPEEDI to single-cell multiome data

SPEEDI can also be used for scATAC-seq and sc multiome datasets. For processing scATAC-seq data, SPEEDI first uses a standard ArchR workflow to process input data (see **STAR Methods**).^10^ Sample batch labels are subsequently inferred, followed by integration and cell type annotation. If sample-paired or true multiome scRNA-seq data are also available, SPEEDI can process both data types.

To demonstrate the application of SPEEDI to single-cell multiome datasets, we processed same cell scATAC and scRNA true multiome data we generated from 14 wild-type female murine pituitary tissue samples (**GSE244132**). SPEEDI identified a total of 60,906 cells meeting QC thresholds. Because SPEEDI independently processes the scATAC-seq and scRNA-seq components of the multiome data, comparison of the cell type annotation of each cell obtained using each data type provides an assessment of the reliability of the automated processing pipeline and batch inference method. The batch labeling was consistent across data types (**Table S3**), indicating SPEEDI captured the technical variation between samples regardless of the type of data. The cell type annotations obtained using the two data modalities were highly consistent (**Figure 4A**). Annotation of individual cells by cell type for the RNA-seq and for the ATAC-seq data after using the SPEEDI pipeline showed a median cell subtype identification overlap of 0.96 as well as a median adjusted Rand index (ARI) of 0.85 (**Figure 4B**). These results support the applicability of the SPEEDI batch inference method and automated pipeline to single-cell ATAC-seq and to single-cell multiome datasets.

**Figure 4.**
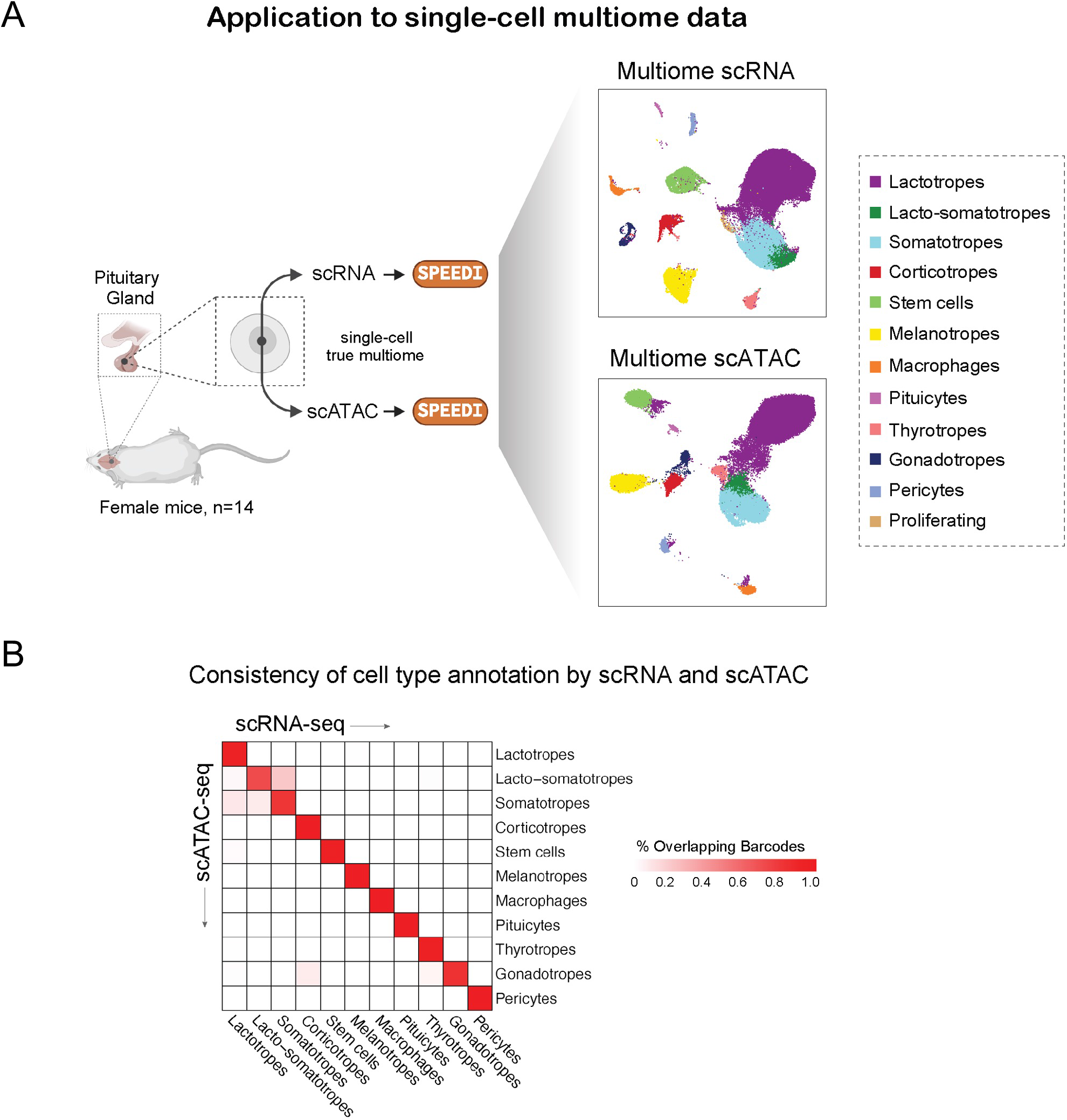
SPEEDI batch inference and framework are highly robust on multiome mouse data. (See also Table S3) (**A**) Same-cell scRNA-seq and scATAC-seq multiome datasets were generated from 14 wild-type female murine pituitaries. Data from each assay were integrated and annotated by the SPEEDI framework. (**B**) Heatmap representation of the contingency table that compares the annotation of individual cells by cell type for the scRNA-seq and for the scATAC-seq data after using the SPEEDI pipeline. The rows represent the cell type annotation for scATAC-seq, and the columns represent those for scRNA-seq. Each cell ranges between 0 and 1, where the value indicates the percentage of overlapping barcodes (median cell subtype identification overlap = 0.96).

### SPEEDI downstream analysis of SARS-CoV-2 single-cell RNA-seq data

To demonstrate the utility of the downstream analysis tools in SPEEDI, we evaluated human PBMC scRNA-seq data from a longitudinal SARS-CoV-2 study.^11^ We used SPEEDI to integrate data from 4 severe seropositive COVID subjects and 5 healthy controls (9 samples in total, **Figure S6A**) and then performed preliminary downstream analyses, including differential expression analysis and functional module discovery in the major immune cell types. The authors used bulk transcriptome data to note downregulation of chemokine CCL3 and cytokine IL1B and upregulation of KLF6 in COVID subjects. After processing the scRNA-seq data through SPEEDI, we applied the differential expression analysis option provided with SPEEDI to compare severe COVID subjects and healthy controls within each cell type. This analysis confirmed the regulatory changes reported in the published bulk RNA-seq analysis and also showed that CCL3 and IL1B are downregulated in CD14 monocytes and that KLF6 is upregulated in CD4 TCM, CD8 Naive, CD8 TEM, B naive, MAIT, and pDC cells (**Table S4**). Furthermore, use of the functional module discovery tool (see Methods) on upregulated and downregulated genes from different cell types revealed additional insights into the specific biological processes associated with severe SARS-CoV-2 infection. For example, upregulated genes in NK cells were found to be associated with positive regulation of the canonical Wnt signaling pathway,^12^ positive regulation of lymphocyte activation,^13^ and the ERAD pathway (**Figure S6B**).^14^ Thus SPEEDI can provide useful automated analysis for single-cell datasets and, even when applied to data that has been previously analyzed, has the potential to facilitate addressing specific questions of interest to the user (in this case cell type-level transcriptome and pathway effects) not previously answered in a published analysis.

## Discussion

As heterogeneous single-cell datasets are generated, appropriate methods for data integration allows robust annotation of cell type and interpretation of the underlying biology.^15^ While many of the currently available integration approaches rely on a pre-defined set of batches for computation,^16^ we demonstrate that subjective batch selection can severely comprise the robustness of study conclusions. SPEEDI addresses this issue by introducing an automated data-driven batch inference method based on information-theoretic principles, therefore overcoming the problem of unknown batch effects.

In addition, we demonstrate that SPEEDI overcomes the challenges of robustness, accessibility, and reproducibility in single-cell data analysis that the biological research community faces. Despite prior efforts to streamline single-cell analysis, current packages either assume user familiarity with coding,^17,18^ or require users to optimize parameter selection.^19,20^ In contrast, SPEEDI’s full automation obviates the requirement of users without bioinformatic background to manually assemble a pipeline, reducing the subjectivity and increasing the reproducibility of analyses. Furthermore, we show that SPEEDI allows for the interrogation of single-cell multi-omics, which could provide a broader and a more profound impact on molecular cellular biology than the traditional single-modality scRNA-seq technology.^21^

In conclusion, SPEEDI provides a streamlined user-friendly processing framework for integration of single-cell omics and multiome datasets. SPEEDI’s one-step pipeline eliminates user parameter setting thereby reducing the need for computational expertise for integration of these valuable datasets improving robustness and reproducibility of subsequent analyses. We show that the data-driven batch labeling method we developed and integrated into SPEEDI improves dataset integration results, even when batch metadata information is available. We offer SPEEDI both as an interactive web application (https://speedi.princeton.edu/) for general users and as an R package. SPEEDI with its data-driven batch labeling method should also be useful for sophisticated bioinformaticians and is easily customizable to allow incorporation of different standalone integration methods. The SPEEDI framework should help a wide range of researchers leverage the power of single-cell omics datasets to provide important new biological insights.

## Acknowledgements

We thank the Single-cell and Spatial Technologies team at the Center for Advanced Genomics Technology, Department of Genetics and Genomic Sciences, the Icahn School of Medicine at Mount Sinai for providing the experimental, computational, data resources, and staff expertise. We used BioRender.com to create the schematic illustrations in Figures 2 and 5. This work was supported by the Defense Advanced Research Projects Agency contract no. N6600119C4022 (S.C.S.), National Institutes of Health grant no. R01DK46943 (S.C.S.) R01GM071966 (O.G.T.), and Simons Foundation grant no. 395506 (O.G.T.).

## Author contributions

Y.W, X.C., E.Z., S.C.S., and O.G.T. conceptualized the study. Y.W. designed and implemented the computational framework, conducted benchmarks and case studies, and wrote the code. W.T. wrote the code and developed the R package, the GitHub repository, and the vignette. W.T. and A.T. set up the web server. F.R.Z. generated the true multiome wild-type mouse pituitary dataset. S.C.S., O.G.T., Y.W., X.C., W.T., A.C., and E.Z. wrote the original draft of the manuscript. All authors proofread the submitted version.

## STAR Methods

### Resource availability

#### Lead contact

Further information and requests for resources should be directed to and will be fulfilled by the lead contact, Olga G. Troyanskaya.

#### Materials availability

This study did not generate new materials.

#### Data and code availability

Human healthy lung scRNA-seq data are available via GitHub (https://github.com/theislab/scib-reproducibility). Peripheral blood mononuclear cell (PBMC) scRNA-seq data from the *S. aureus* infection study are publicly available on GEO (GSE220190). Sc multiome wild-type mouse pituitary data are available on GEO (GSE244132). PBMC scRNA-seq data from the longitudinal SARS-CoV-2 study are publicly available on GEO (GSE206283). To review GEO accession GSE244132 currently as a reviewer or editor, please go to https://www.ncbi.nlm.nih.gov/geo/query/acc.cgi?acc=GSE244132 Enter token **gxmbigygrvslpch** into the box.

The source code of SPEEDI is available as an R package on GitHub at https://github.com/FunctionLab/SPEEDI/. The SPEEDI web server is publicly available at https://speedi.princeton.edu/.

Any additional information required to reanalyze the data reported in this paper is available from the lead contact upon request.

### Methods details

#### Input format and external data

For scRNA-seq data, SPEEDI accepts raw data in unique molecular identifier (UMI) count matrix format. For scRNA, sample data should be filtered data generated by Cell Ranger in the MEX format. SPEEDI also works with H5 format files. If working with MEX files, sample data should always follow the standard naming convention (three files with names barcodes.tsv.gz, features.tsv.gz, and matrix.mtx.gz). If working with H5 files, sample data file names should always end with filtered_feature_bc_matrix.h5. For scATAC-seq data, SPEEDI requires the fragment files instead. For all data types, SPEEDI utilizes “hg38” and “mm10” as the reference genome annotation for human and mouse, respectively. More specifically, “hg38” represents the UCSC full genome sequence hg38 for Homo Sapiens available in Bioconductor package “BSgenome.Hsapiens.UCSC.hg38”, while “mm10” comes from the “BSgenome.Mmusculus.UCSC.mm10” package, which is the UCSC full genome sequences for Mus musculus version mm10 based on GRCm38.p6. If the experiment includes multiple conditions, SPEEDI also optionally accepts a metadata file specifying the sample IDs and the experimental condition of each sample for downstream differential analysis.

#### Parameter optimization for single-cell RNA-seq quality control

The standard practice for single-cell RNA-seq data quality control (QC) is to perform cell filtering based on at least two covariates: the number of detected transcripts per barcode, and the number of mitochondrial genes per barcode.^22^ Experience shows that additional covariates should also be considered, including but not limited to the number of ribosomal RNA molecules (both large and small subunits) per barcode as well as number of hemoglobins per barcode,^23^ which typically appears in PBMC (peripheral blood mononuclear cell) datasets as contamination. Accordingly, for scRNA-seq we implemented four QC filters: transcripts per barcode, mitochondrial genes per barcode, ribosomal RNA molecules per barcode and hemoglobin per barcode, with the last affecting mostly PBMC analysis.

Current practice of single-cell QC is to manually inspect the distribution of the aforementioned covariates per barcode and hand pick a dataset-specific outlier threshold. However, this approach compromises reproducibility as a threshold too loose or too stringent could potentially lead to different conclusions.

To parametrize the QC process in a data-driven fashion, SPEEDI utilizes the Kneedle algorithm that systematically finds the elbow or knee point in the distribution of transcript counts per cell.^24^ While the term “elbow” is typically applied to the inflection of a convex curve and “knee” to that of a concave curve, for simplicity here we use “elbow” for both situations.

Conventionally, the elbow point is considered as the turning point where a distribution changes its behavior due to some latent factor. In single-cell data analysis, it can be interpreted as the point where the profiled barcodes transit into a different quality phase. In the case of a unimodal distribution, there are typically left and right elbow points that represent the lower and the upper threshold, respectively, for our algorithm. Data below this lower threshold are normally associated with broken cells where cytoplasmic mRNAs are escaping from the cell membrane.

To calculate the lower threshold of UMI counts *L*_*UMI*_, SPEEDI begins by creating a 100-group histogram of the UMI count covariate. The histogram effectively creates a set of discrete data points *D*_*i*_, where

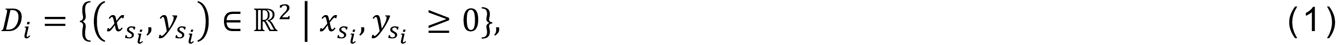

*x*_*i*_ is the *i*^*th*^ percentile of UMI counts where *i* ∈ ℕ^+^ and 0 < *i* ≤ 100;

*y*_*i*_ is the frequency of the *i*^*th*^ percentile of UMI counts and *y*_*i*_ ∈ ℕ^+^.

To find the left elbow point from the curve formed by point set *D*_*i*_, SPEEDI first finds the subset 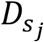 between points 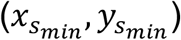 and 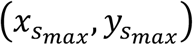, such that

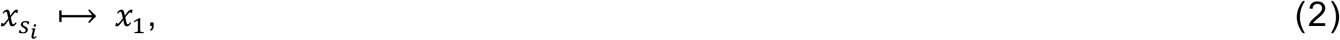

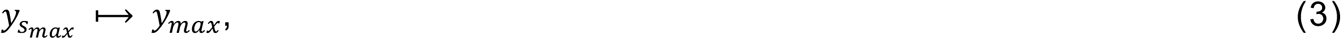

The Kneedle algorithm first rotates the curve segment defined by 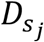 clockwise for *θ* degrees so that the curve is concave and the line formed between 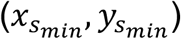 and 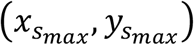 is horizontal. It then finds the local maximum 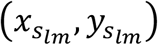 of the rotated curve. The lower elbow point is thus defined as

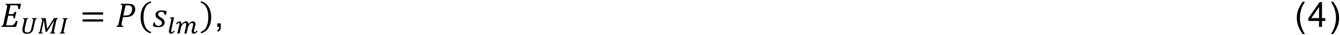

where *P*(*s*_*lm*_) is ^the^ *s*_*lm*_^*th*^ percentile of UMI counts, 0 < *s*_*lm*_ < 100.

To restrict the algorithm from filtering too many cells, SPEEDI conservatively imposes an upper limit of 1,000 UMI counts, such that the final threshold of gene count is defined as

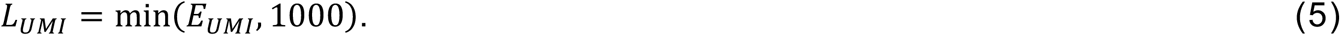

Apart from low quality cells, SPEEDI also filters cells that have excessively high UMI counts. SPEEDI defines the upper limit to be

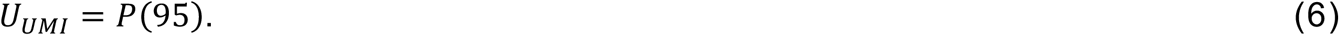

Similarly, let *E*_*MT*_ be the right elbow point calculated from the curve defined by the distribution of mitochondrial reads. We define the upper limit of mitochondrial content in percentage as

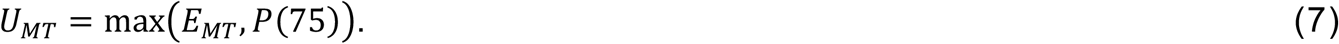

For additional covariates, we uniformly choose the 99^*th*^-percentile as the upper threshold. Note that SPEEDI performs QC on individual sample datasets to avoid detecting false positive outliers that are distributed differently because of biological or technological factors (e.g., rare cell types detected in a subset of samples or different sequencing depths). Cell cycle scoring is also performed by calling the “CellCycleScoring” function with canonical cell cycle markers,

After QC, the framework merges the processed samples into one dataset for normalization. SPEEDI uses Seurat’s “SCTransform” function with v2 regularization to normalize the merged count matrix.^25^ All QC metrics as well as cell cycle scores are used as regression covariates in normalization. The normalized matrix is then scaled and centered.

#### Single-cell ATAC-seq quality control

SPEEDI processes sample fragment files with ArchR.^10^ Each input read count by cell matrix is converted to a tile matrix by binning the genome-wide fragments into tiles of 500 bps. SPEEDI selects scATAC-seq quality cells with fixed thresholds. Cells with a number of fragments between 3,000 and 30,000, TSS enrichment > 12, and nucleosome ratio < 2 are retained. Doublets are removed using the “filterDoublets” function of ArchR under default settings.

#### Dimension reduction

After normalization, SPEEDI projects the higher dimensional matrix into a lower dimensional orthogonal embedding of cells depending on the appropriate data type. For scRNA-seq, PCA (principal component analysis) is performed on the normalized expression matrix. For scATAC-seq, LSI (latent semantic indexing) is performed on the tile matrix using the ArchR function “addIterativeLSI” with settings iterations = 2 and varFeatures = 20000.^26^

#### Batch inference

The batch inference method uses the unique structure of single cell data (containing both different samples as well as groups of different cell types within each sample) as the basis for an information theoretic method to identify data-defined batches, whatever their cause. At a high level, the method is based on the idea that data from the same cell type in different samples should be more similar than different cell types within each sample.

The sample batch inference algorithm makes the following assumptions:

1. Let the input normalized matrix be 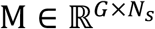 where element 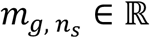 is the normalized of the *g*^*th*^ gene in the *n*^*th*^ cell of the *s*^*th*^ sample. SPEEDI requires |*G*| > 30.
2. Within each sample, the separation among the cell types is preserved.
3. Any sample can belong to any batch and a sample can be its own unique batch.
4. The cell types contained within each sample do not have batch variations relative to each other.
5. Batch effects do not intermix cells of distinct types or states from different samples. In other words, the expression profiles of cells of cell type A in sample *x* are more correlated to cells of cell type A sample *y* than to cells of cell type B or C in sample *y*.
6. Samples do not have to contain identical cell types.

The process begins with an initial round of clustering at a low resolution that clusters similar cells from different samples. These clusters represent a higher-level cell ontology similar to a cell type or state. This allows identification of batches on a per-cell-type basis, maintaining its sensitivity to local batch effects where only some cell types are affected by batch effects. SPEEDI uses a grid-search approach to find the optimal clustering resolution, using a search space ranging from 0.05 to 1 with 0.01 increments. We define the optimal clustering resolution as being the resolution that produces the highest average Silhouette width (ASW) of all samples.^27^

The SPEEDI batch labeling method then iteratively finds differentially distributed groups of samples within clusters that contain at least 1% of all cells using Hill numbers *D*^q^, an information-theoretic metric traditionally used to measure biological diversity.^28^

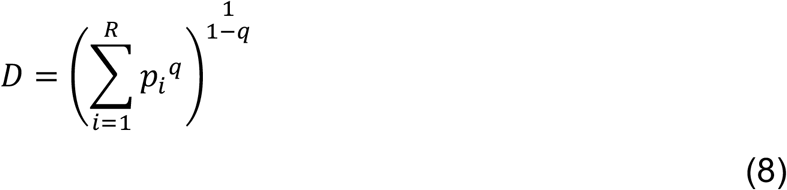

When q = 2, the formula becomes the Inverse Simpson Index (ISI). Korsunsky et al. implemented a variant to ISI by adding a distance-based weight to the original index, namely Local Inverse Simpson’s Index (LISI).^4^ Therefore, for each cluster, SPEEDI first calculates an LISI score that quantifies sample distributions. The unscaled LISI score measures the effective number of batches given observations and ranges from 0 to total number of samples within the target cluster. However, given that the LISI metric is influenced by cell population size diversity, the original implementations of LISI are inappropriate for SPEEDI. Therefore, the SPEEDI algorithm introduces a penalized calculation of sample-level LISI (sLISI) scores using the frequency of occurrence of cells in each sample as weights. Finally, the sLISI scores are scaled such that a score closer to 1 indicates sample well-mixing within the given cluster, while a score closer to 0 means the sample is locally enriched. As a result, samples with significantly small sLISI scores form putative batches. To detect whether a group of low-scoring samples are outliers, SPEEDI employs Dixon’s Q-test. Given ordered sLISI scores

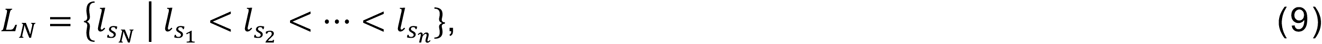

where 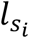 is the sLISI score for sample *s*_*i*_, SPEEDI calculates the difference between each score and its subsequent neighbor, {*d*_*N*_}, where

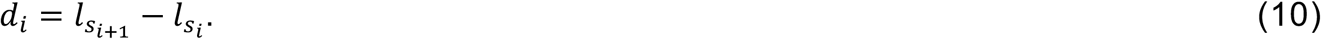

The goal is to find the outlier in *D*_2_ with Dixon’s Q-test, which represents the most statistically significant (P<0.05) gap between two consecutive scores 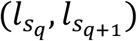 in *L*_N_ such that samples {*s*_1_, …, *s*_q_} are considered to be one group, and samples {*s*_q+1_, …, *s*_*n*_} form another group. SPEEDI iteratively assigns batch labels with the outlier detection method to ensure that all significant differences between each score are accounted for.

#### Batch integration

The framework returns a finalized batch assignment for all samples. SPEEDI uses the inferred batch covariate to subset the merged multi-sample dataset into a list of batch datasets. The default integration for scRNA-seq in SPEEDI is to run the Seurat RPCA method by calling the “FindIntegrationAnchors” function followed by the “IntegrateData” function. For scATAC-seq data, if SPEEDI detects that batch effects are present, the “addHarmony” function will then be called to integrate the lower embedding of each sample with SPEEDI-inferred batches.

#### Cell type annotation

After samples are integrated into a batch-corrected matrix, SPEEDI annotates cells with a reference-based approach. For scRNA-seq, the default version of SPEEDI implements the Seurat label transfer algorithm that projects an identity to each query cell using some prior annotated reference data by calling the “FindTransferAnchors” function. Since ArchR also adopts Seurat label transfer, the same algorithm also applies to scATAC-seq data. Clustering is then performed with the “addClusters” function, and cell types are projected from a reference gene expression dataset onto the query chromatin accessibility dataset using the ArchR function “addGeneIntegrationMatrix”.

SPEEDI can be easily incorporated into other single-cell annotation tools such as SingleR.^7^ On top of per-cell label transferring, SPEEDI introduces a majority-voting regime that further refines cell labeling. The majority-voting regime begins with an over-clustering of the integrated dataset at a higher resolution, where each cluster represents a cell phenotype that serves as a surrogate to a parent cell type or state. It then loops through each cluster and finds the most represented predicted cell types and states among cells within the query cluster. Since the data is over-clustered, this process eventually merges clusters representing the same cell identities together for a data-driven cell label annotation landscape.

#### Evaluation metrics

Leucken et al. defined two categories of metrics to evaluate integration results.^5^ The first category measures the conservation of biological variance. The second category reflects the removal of batch effects. In this study, a total of four appropriate metrics are selected as evaluation criteria. Metrics from the first category consider how well cells of different identities are separated for each sample. Metrics from the second category measure the degree of overlapping of cells from each sample for each cell type.

##### Per-sample ASW

The average silhouette width (ASW) is a classical metric that evaluates the validity of a clustering partition based on between-cluster proximities.^27^ In single-cell data science, a clustering of cells can be represented by labels such as sample IDs, sequencing protocols, cell type identities, etc. In the original definition, the ASW ranges between -1 and 1 such that -1 means strong overlapping and 1 means perfect separation between clusters.

The per-sample ASW score for cell type separation is computed based on sample IDs and scaled between 0 and 1 using Equation (11) for each sample. A score closer to 1 represents well-conserved cell types.

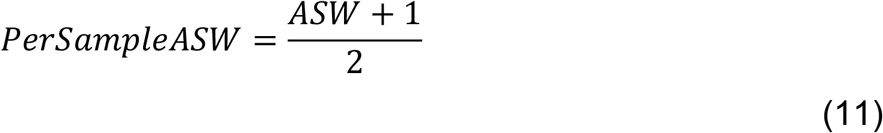

##### Per-sample graph connectivity

The graph connectivity metric quantifies the average connectivity of the subgraph of cells in target cell type *c* ∈ *C* in comparison to the largest connected component (LCC) for all cells of all types in a kNN graph. A score of 1 indicates that strong local cell type coherence is preserved post integration. A score is computed for each sample using Equation (12).

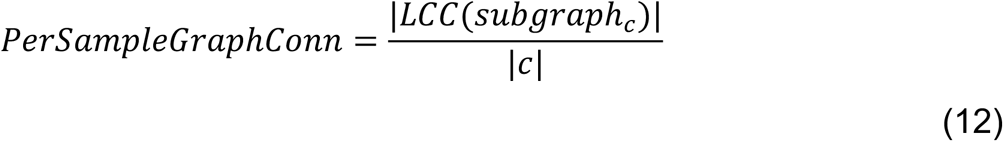

##### Per cell-type GraphConn

The per cell-type graph connectivity metric is defined similarly to the per-sample GraphConn metric, except for that it now averages the connectivity of all sample *s* ∈ *S*. A score of 1 indicates that all cells of the same type are strongly connected post integration. A score is computed for each cell type using Equation (13).

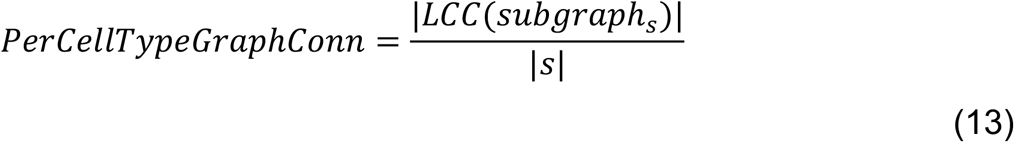

##### Per cell-type graph LISI

The local inverse Simpson’s index (LISI),^4^ as explained in Equation (8), was further adapted by Leucken et al. to use the graph structure of the integrated data to calculate shortest path length as distance between two nodes.^5^ Like the original LISI, the graph LISI ranges from 1 to the total number of batches. The metric is then scaled with Equation (14) to a score between 0 and 1 for each cell type such that 1 indicates perfect cell type preservation.

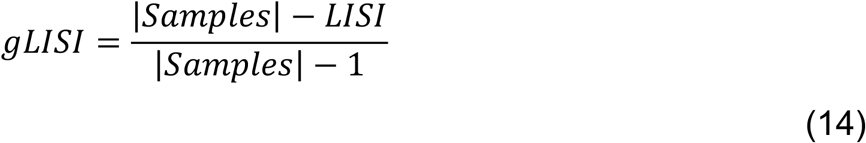

#### Benchmarking studies

The human healthy lung single-cell expression data was originally collected by Vieira Braga et al. which includes three diverse datasets across different sequencing technologies and spatial locations.^8^ Two of the datasets were generated by 10X Chromium and the remaining dataset was generated by Drop-seq. The Drop-seq dataset and one of the 10X datasets contain cells profiled from upper and lower airways which were retrieved from deceased transplant donors. The other 10X dataset consists of parenchyma cells from resected lung tissue. These three datasets were further processed by Luecken et al. to include 16 samples composed of 32,472 cells and 15,148 genes, where a re-annotation of cells was performed since 10X and Drop-seq datasets used different annotation terms.^5^

The raw count expression matrix was downloaded from Luecken et al. and was re-normalized using SPEEDI protocol. Batches were labeled for benchmarking using either the sample ID, the dataset ID, or the SPEEDI data-driven method. P-values were calculated from the paired two-sample non-parametric Wilcoxon rank sum test and Bonferroni-adjusted.

#### Human bloodstream *S. aureus* infection study

The scRNA-seq human *S. aureus* infection PBMC dataset was generated by our lab and previously reported in Chen et al.^9^ 20 adult patients with culture-confirmed infection were selected with 10 MRSA patients and 10 MSSA patients. The sequencing batch labels were reported in an accompanying metadata file. We extracted the H5 files from 10x Genomics Cell Ranger v3.1.0 and reprocessed the raw count matrices using SPEEDI. For sample integration, we used two sets of batch labels: (a) the sample ID labels, and (b) SPEEDI-inferred batch labels. Two different integrated datasets were subsequently obtained based on this input batch covariate. To assign cell types, we obtained a reference dataset by extracting data from 21 uninfected healthy individuals that were also reported and curated in Chen et al. For illustration purposes, we focused the analysis on a subset of cells that are T/NK leukocytes.

#### Female mouse pituitary study

##### Animal and pituitary collection

The pituitaries used in this study were collected from wild-type untreated female C57BL/6 mice aged 11 weeks. All murine sample collection was conducted at McGill University (Montreal, Quebec, Canada). All animal experiments were performed in accordance with institutional and federal guidelines and were approved by the McGill University and Goodman Cancer Centre Facility Animal Care Committee DOW-A (Protocol 5204). We have complied with all ethical regulations and institutional protocols. Once dissected, the pituitaries were individually collected, immediately snap-frozen, and stored at -80C until the assay was started.

##### Nuclei isolation from the murine pituitary

Nuclei isolation was performed as described in previous literature.^29,30^ Briefly, each snap-frozen pituitary was thawed individually on ice. RNAse inhibitor (NEB MO314L) was added to the homogenization buffer (0.32 M sucrose, 1 mM EDTA, 10 mM Tris-HCl, pH 7.4, 5mM CaCl2, 3mM Mg(Ac)2, 0.1% IGEPAL CA-630), 50% OptiPrep (Stock is 60% Media from Sigma; cat# D1556), 35% OptiPrep and 30% OptiPrep right before isolation. Each pituitary was homogenized in a dounce glass homogenizer (1ml, VWR cat# 71000-514), and the homogenate filtered through a 40 mm cell strainer. An equal volume of 50% OptiPrep was added, and the gradient centrifuged (SW41 rotor at 9200rpm; 4C; 25min). Nuclei were collected from the interphase, washed, resuspended in 1X nuclei dilution buffer (10X Genomics), and counted (Nexcelom K2 counter).

##### True (sn) multiome assay

True (sn) multiome was performed following the Chromium Single Cell Multiome ATAC and Gene Expression Reagent Kits V1 User Guide (10x Genomics, Pleasanton, CA). Nuclei were counted as described above, transposition was performed in 10 ml at 37C for 60 min targeting 10,000 nuclei, before loading of the Chromium Chip J (PN-2000264) for GEM generation and barcoding. Following post-GEM cleanup, the library was pre-amplified by PCR, after which the sample was split into three parts: one part for generating the snRNA-seq library, one part for the snATAC-seq library, and the rest was kept at -20C. snATAC and snRNA libraries were indexed for multiplexing (Chromium i7 Sample Index N, Set A kit PN-3000262, and Chromium i7 Sample Index TT, Set A kit PN-3000431 respectively).

##### Quality control (QC) and sequencing of libraries

The libraries were quantified by Qubit 3 fluorometer (Invitrogen), and quality was assessed by Bioanalyzer (Agilent). The libraries were sequenced first in a MiSeq (Illumina) to assess the reads and balance the sequencing pools and then sequenced in a NovaSeq 6000 (Illumina) at the New York Genome Center (NYGC) following 10X Genomics recommendations.

#### Optional downstream analyses

To run downstream analyses, the user is required to provide a metadata table where row names are sample names and column names are metadata attributes of interest. Note that each metadata attribute should contain exactly two unique labels (e.g., “disease” and “control”). SPEEDI has two different downstream analysis options. First, SPEEDI can run cell-type specific differential expression analyses. For each cell type, SPEEDI uses the Wilcoxon Rank-Sum test (via Seurat’s “FindMarkers” function) to perform differential expression using the sample labels associated with a given metadata attribute. Results are filtered using an adjusted p-value threshold of 0.05, a log fold change threshold of 0.1, and a min.pct threshold of 0.1 (genes must be expressed in at least 10% of cells in one of the two groups associated with the metadata attribute). Next, pseudobulk analysis is performed using DESeq2.^31^ First, pseudobulk counts are calculated for each cell type and DESeq2 is used to find differentially expressed genes (via Wald test). Results are then filtered using a p-value of 0.05. Genes that pass both single-cell and pseudobulk differential analysis constitute the final list of differentially expressed genes. Differential expression results for each metadata attribute are written to a TSV file.

SPEEDI can also perform functional module discovery (FMD) to find functionally connected gene clusters using HumanBase.^32^ By default, SPEEDI uses the lists of genes generated by the cell-type specific differential expression analysis described above. For each metadata attribute and cell type, the list of genes is first divided into positive fold change and negative fold change subsets. If at least 20 genes remain after fold change filtering, SPEEDI runs FMD. Results are written to a CSV file and include a URL that allows users to see their full results as well as a table that contains gene ontology (GO) enrichment results. Note that FMD can only be run with human genes.

## Supplemental information titles and legends

### Supplemental figures titles and legends

**Figure S1. Strong batch effects are present in the human PBMC scRNA-seq datasets across all cell types before integration. (Related to Figure 3)**

(**A**) UMAP shows cells are segregated by sample IDs indicating strong batch effects present in all cell types of the human PBMC scRNA-seq data.

(**B**) Louvain clustering at resolution of 0.74 produced 47 clusters. Adjusted Rand Index of cells between sample IDs and cell clusters is 0.39.

**Figure S2. Integrated data for all cell types in human PBMC scRNA-seq data colored by sample IDs. (Related to Figure 3)**

(**A**) UMAP of sample ID-based integration.

(**B**) UMAP of data-defined batch-based integration.

**Figure S3. Integrated data for all cell types in human PBMC scRNA-seq data colored by cell types. (Related to Figure 3)**

(**A**) After integrating using sample IDs, the UMAP colored by cell type shows broken cell populations (i.e. *CD14*^+^ Monocytes and Tregs).

(**B**) Broken cell populations are minimal after integrating using SPEEDI data-derived batch labels.

**Figure S4. Sample-based integration fails to recover distinct *CD8***^***+***^ **T cell populations. (Related to Figure 3)**

(**A**) UMAPs with cells predicted to be *CD8*^+^ memory T cells highlighted. Cells predicted to be *CD8*^+^ are spread across the entire NK/T lymphocyte landscape in the sample ID-based integration, whereas those in the data-defined batch-based integration are much more concentrated.

(**B**) UMAPs of *CD8A*^+^ cells indicating the distribution of the *CD8*^+^ T cell subset.

(**C**) UMAPs of *CD3D*^+^ cells indicating the distribution of T cells.

**Figure S5. UMAPs of gene expression of canonical markers confirm SPEEDI inferred cell types. (Related to Figure 3)**

(**A-E**) UMAPs of *NKG7*^+^ cells, *SELL*^+^ cells, *TNFRSF4*^+^ cells, and *FOXP3*^+^ cells.

**Figure S6. Demonstration of SPEEDI downstream analysis capabilities**.

(**A**) UMAP of integrated scRNA-seq data from a longitudinal SARS-CoV-2 study.

(**B**) Functional module discovery on upregulated NK genes.

### Supplemental tables titles

**Table S1. SPEEDI auto calibration of parameters. (Related to Figure 1)**

**Table S2. Summary of cell type representations across all samples in the human lung atlas data compendium. (Related to Figure 2)**

**Table S3. SPEEDI independently infers highly corresponding batch groups in paired scRNA-seq and scATAC-seq from mouse multiome data. (Related to Figure 4)**

**Table S4. Differentially expressed genes from SPEEDI downstream analysis of longitudinal SARS-CoV-2 study**.

