## Supplemental Figures for "Automated single-cell omics end-to-end framework with data-driven batch inference"

**Figure S1. Strong batch effects are present in the human PBMC scRNA-seq datasets across all major cell types before integration. (Related to Figure 3)**

A

B

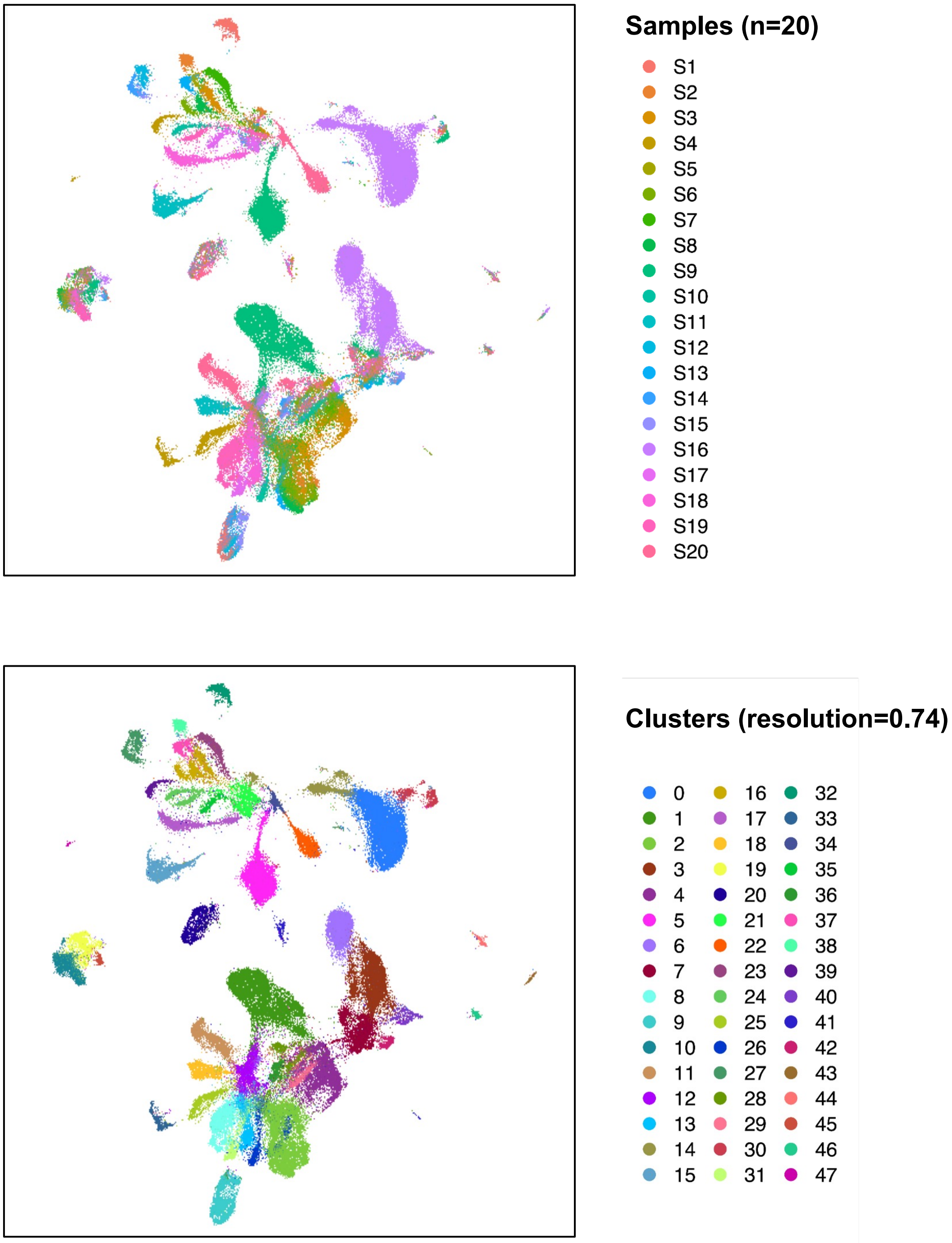

**Figure S2. Integrated data for all cell types in human PBMC scRNA-seq data colored by sample IDs. (Related to Figure 3)**

A

B

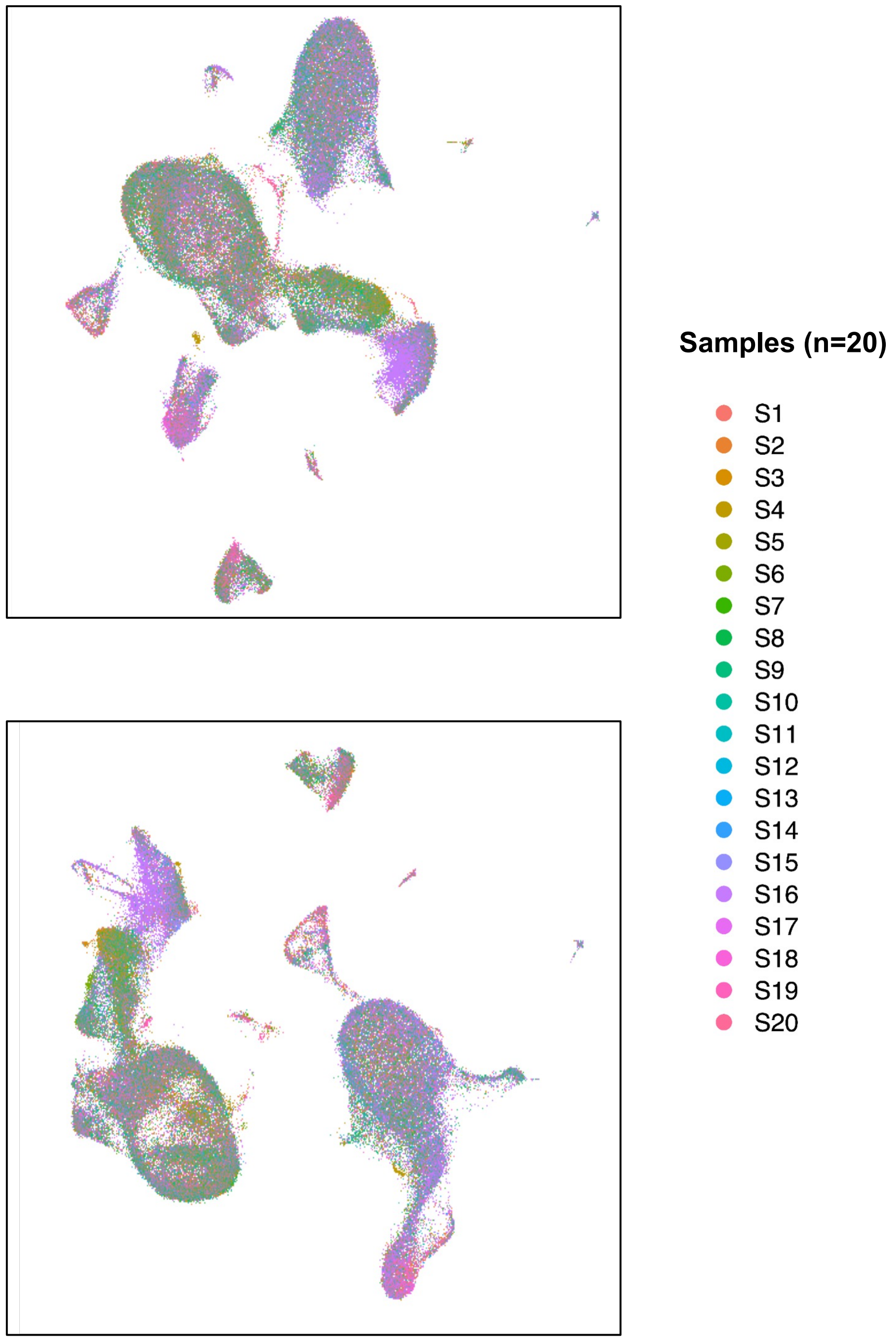

**Figure S3. Integrated data for all cell types in human PBMC scRNA-seq data colored by cell types. (Related to Figure 3)**

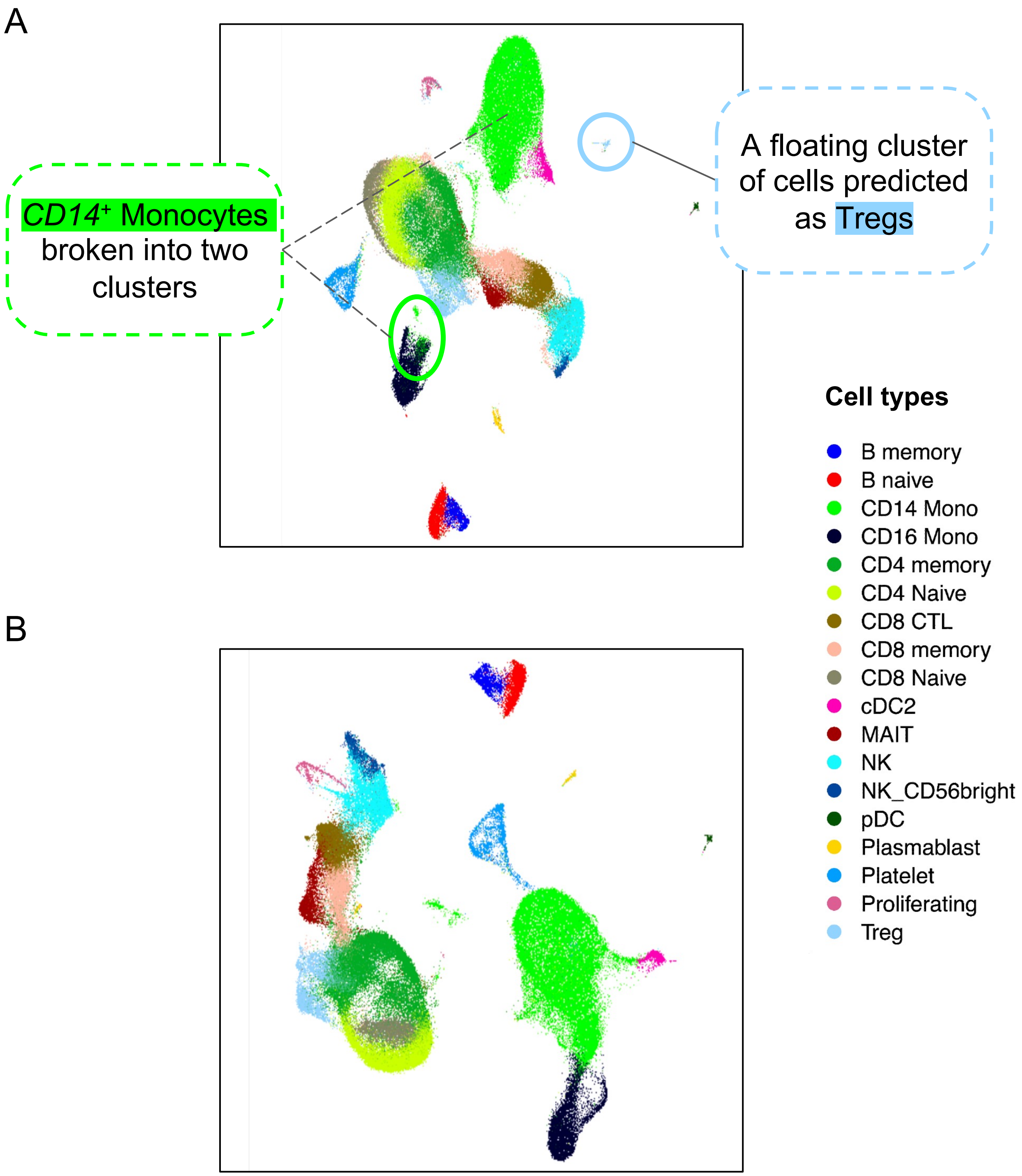

**Figure S4. Sample-based integration fails to recover distinct *CD8⁺* T cell populations. (Related to Figure 3)**

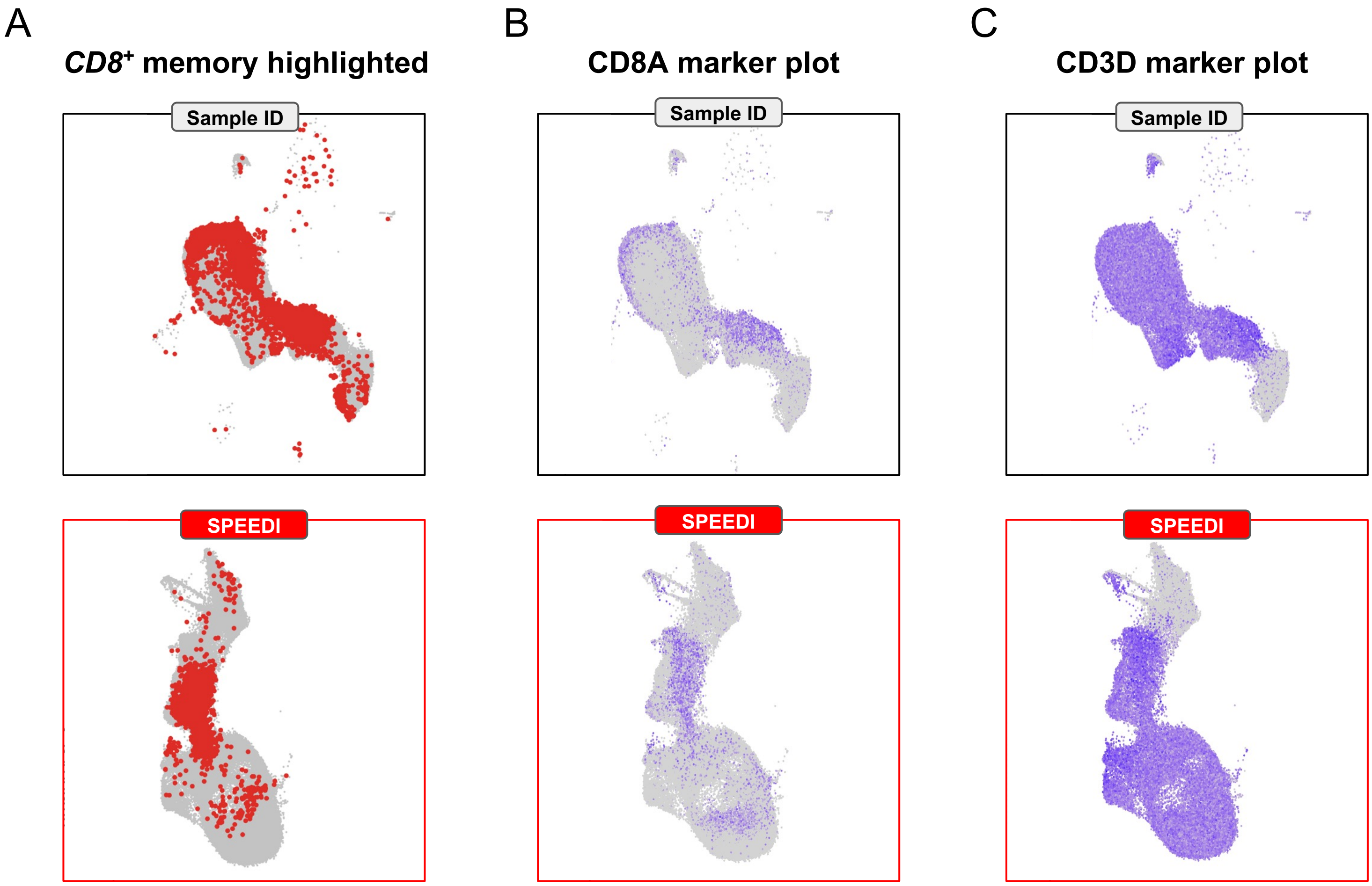

**Figure S5. UMAPs of gene expressions of canonical markers confirm SPEEDI inferred cell types. (Related to Figure 3)**

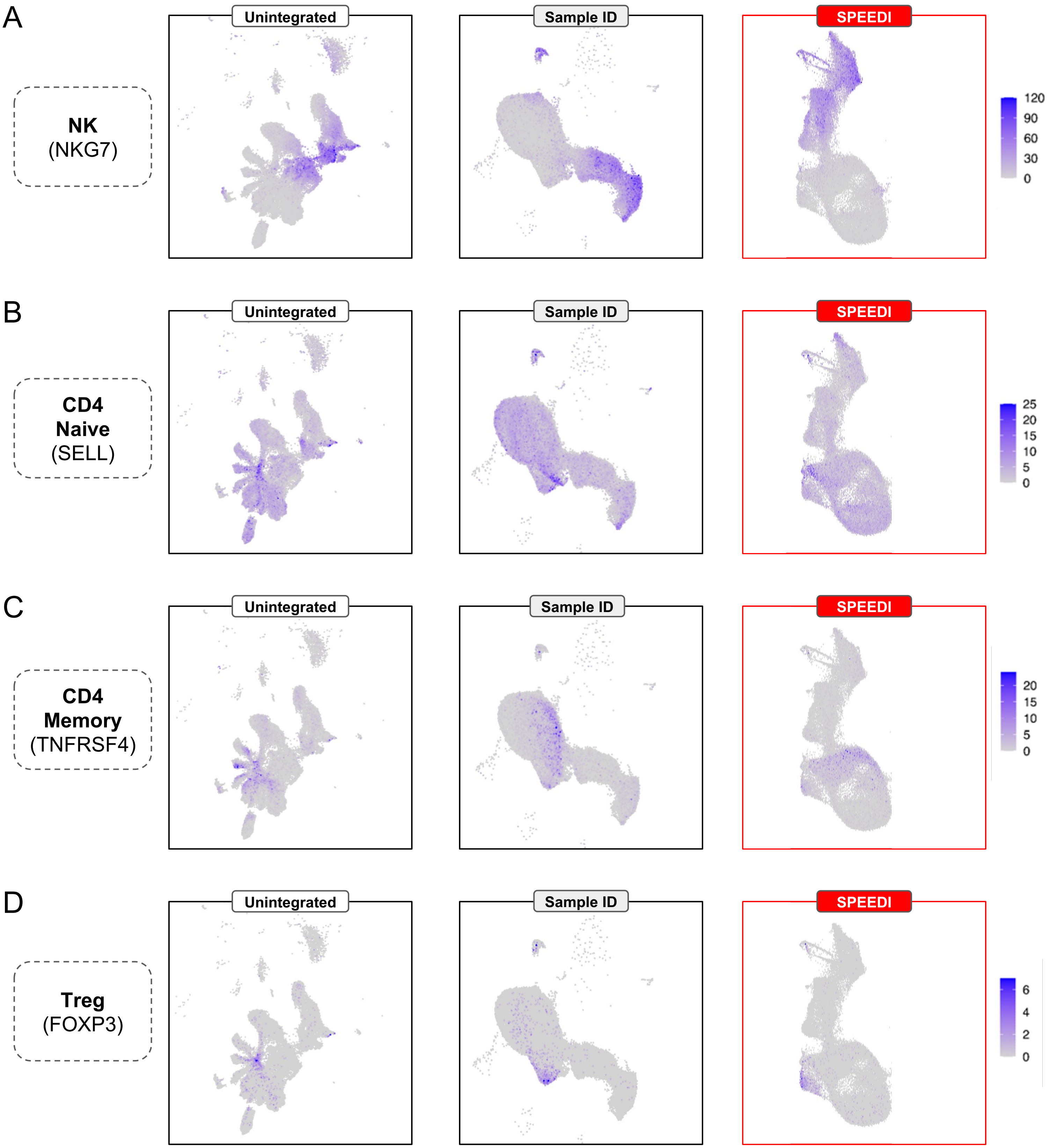

**Figure S6. Demonstration of SPEEDI downstream analysis capabilities.**

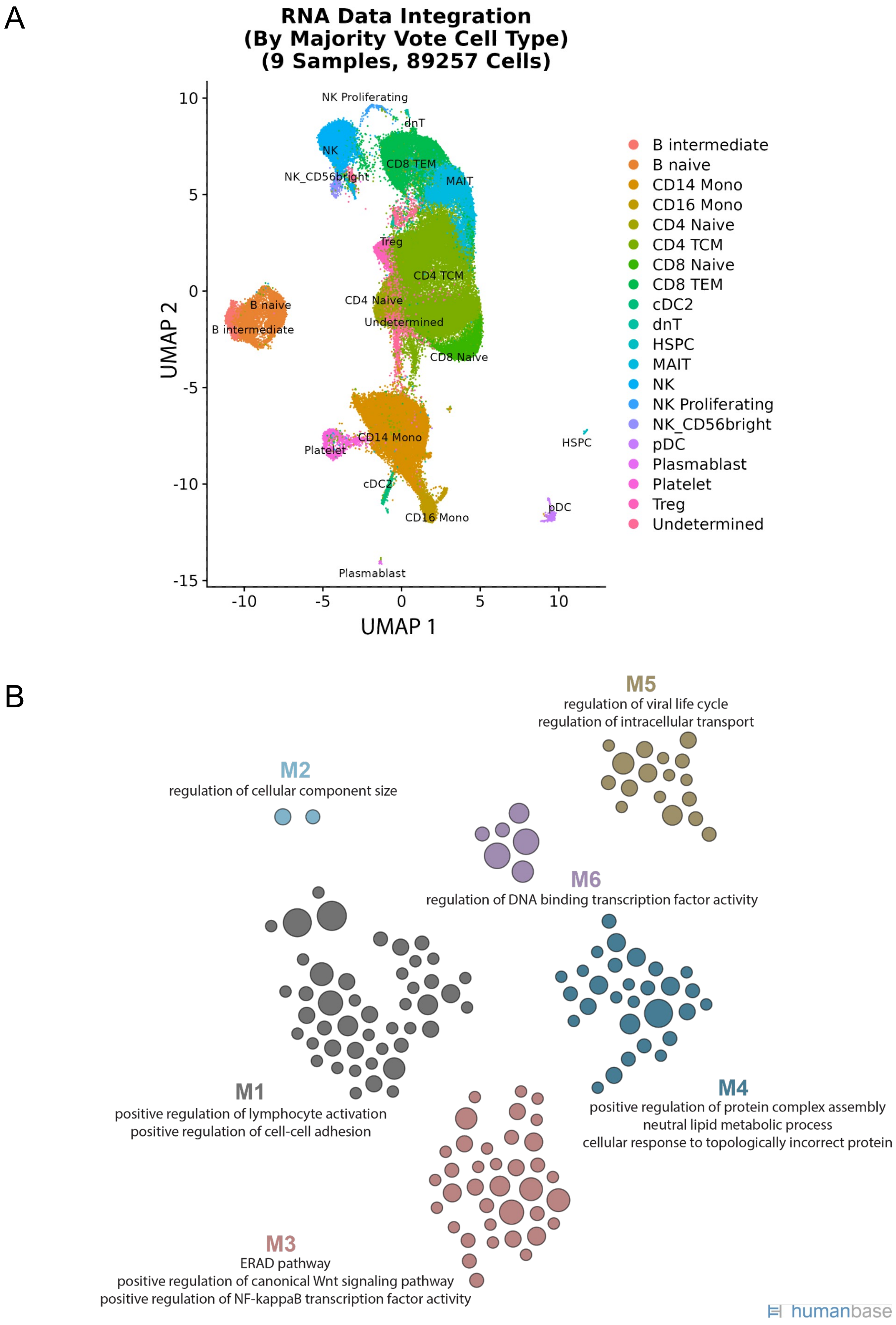
